# SARS-CoV-2 Spike Protein Regulation of Angiotensin Converting Enzyme 2 and Tissue Renin-Angiotensin Systems: Influence of Biologic Sex

**DOI:** 10.1101/2021.09.14.460275

**Authors:** Charles M. Ensor, Yasir AlSiraj, Robin Shoemaker, Jamie Sturgill, Suresh Keshavamurthy, Elizabeth M. Gordon, Brittany E. Dong, Christopher Waters, Lisa A Cassis

## Abstract

Angiotensin converting enzyme 2 (ACE2) is an enzyme that limits activity of the renin-angiotensin system (RAS) and also serves as a receptor for the SARS-CoV-2 Spike (S) protein. Binding of S protein to ACE2 causes internalization which activates local RAS. ACE2 is on the X chromosome and its expression is regulated by sex hormones. In this study, we defined ACE2 mRNA abundance and examined effects of S protein on ACE2 activity and/or angiotensin II (AngII) levels in pivotal tissues (lung, adipose) from male and female mice. In lung, ACE2 mRNA abundance was reduced following gonadectomy (GDX) of male and female mice and was higher in XX than XY mice of the Four Core Genotypes (FCG). Reductions in lung ACE2 mRNA abundance by GDX occurred in XX, but not XY FCG female mice. Lung mRNA abundance of ADAM17 and TMPRSS2, enzymes that shed cell surface ACE2 and facilitate viral cell entry, was reduced by GDX in male but not female mice. For comparison, adipose ACE2 mRNA abundance was higher in female than male mice and higher in XX than XY FCG mice. Adipose ADAM17 mRNA abundance was increased by GDX of male and female mice. S protein reduced ACE2 activity in alveolar type II epithelial cells and 3T3-L1 adipocytes. Administration of S protein to male and female mice increased lung AngII levels and decreased adipose ACE2 activity in male but not female mice. These results demonstrate that sex differences in ACE2 expression levels may impact local RAS following S protein exposures.

## Introduction

The angiotensin converting enzyme 2 (ACE2) full length cDNA was cloned by two independent groups in 2000.(1, 2) A few years later, ACE2 was identified as the functional receptor for the SARS-CoV-1 virus(3), and more recently demonstrated as the major receptor for SARS-CoV-2(4, 5 2020 2020). Over the course of several years since its identification as the receptor for SARS-CoV-1, studies demonstrated that binding of SARS-CoV-1 Spike (S) protein to ACE2 resulted in internalization of cell surface ACE2 (cACE2) as the virus entered cells(6–8), with some studies demonstrating potential recycling of ACE2 to the cell membrane(6) and others indicating loss of available cACE2(8). These findings, while interesting, were not extensively considered in the context of the renin-angiotensin system (RAS) until the pandemic of SARS-CoV-2 coronavirus disease-2019 (COVID-19), with current (August, 2021) global estimates of 216,867,420 confirmed cases and 4,507,837 deaths according to statistics from the World Health Organization (www.who.int). Within the US (August, 2021), there have been 38,666,050 confirmed cases and 632,893 deaths from COVID-19. The magnitude of the pandemic, coupled with significant death of the global population, demonstrate the need for understanding mechanisms that impact the severity of COVID-19 and its post-acute sequalae.

Amongst several risk factors for the severity of COVID-19, there appear to be consistent data indicating that male sex is associated with increased hospitalizations (12 males for every 10 females), ICU admissions (18 males for every 10 females), and deaths (13 males for every 10 females) (see https://globalhealth5050.org/the-sex-gender-and-covid-19-project/). These data are interesting as confirmed global cases of COVID-19 appear to be similar between males and females (11 males for every 10 females). Thus, despite a similar rate of SARS-CoV-2 infection between males and females, males infected with the virus have a worse prognosis.

In the context of the RAS, the receptor for SARS-CoV-2, ACE2, limits activity of local systems by converting the vasoconstrictor angiotensin II (AngII) to the vasodilator angiotensin-(1-7) (Ang-(1-7)).(1, 2, 9, 10) This enzyme function of ACE2, which takes place within the ectodomain portion of the protein that also houses the SARs-CoV-2 S protein receptor binding domain (RBD), limits activity of local RAS systems in cell types relevant to the cardiovascular system (e.g., cardiac myocytes, kidney tubular or mesangial cells, adipocytes, brain neurons). Thus, as has been suggested in several reviews, if cACE2 is internalized by SARS-CoV-2 S protein binding and viral entry, potential stimulation of local RAS activity would predictably augment AngII-mediated effects that influence the severity of COVID-19 (i.e., loss of lung barrier function).

An additional interesting feature of ACE2 and SARS-CoV-2 is the influence of certain proteases to cleave the ectodomain of cACE2 while also having an impact on viral cell entry. Specifically, ADAM-17 is a metalloproteinase that cleaves a soluble form of ACE2 (sACE2) and has also been implicated in SARs-CoV-2 S protein entry(11, 12). Similarly, the serine protease TMPRSS2 is essential in cleavage of SARS-CoV-2 S proteins and viral cell entry and also capable of cleaving the ectodomain of cACE2.(11, 13) Release of sACE2 by these enzymes has been predicted to activate the local RAS by limiting the ability of cACE2 to catabolize locally formed AngII.(14) Given these features of ACE2 to influence both local RAS system activity and also serve as the conduit for SARS-CoV-2 viral cell entry, it is important to identify regulatory mechanisms of ACE2 that might influence COVID-19 severity.

ACE2 is located on the X chromosome.(15) While most genes on the X chromosome do not exhibit gene dosage effects because of the process of X chromosome inactivation (XCI), it is unclear if ACE2 escapes this random process in various cell types and exhibits gene dosage effects in XX females. In addition to its X chromosome location, ACE2 has also been demonstrated to be under the influence of sex hormone regulation, with several studies indicating positive effects of estrogen to stimulate ACE2 gene transcription.(16–20) Taken together, gene dosage effects from two X chromosomes, coupled with estrogen stimulation of ACE2 gene transcription would potentially result in higher levels of residual cACE2 in pivotal cell types of females than males following exposure to SARS-CoV-2. In this study, we hypothesized that sex defining factors regulate residual cACE2 in pivotal cell types following exposure to SARS-CoV-2 S protein, resulting in a more stimulated local RAS and augmented severity of males than females. To test this hypothesis, we investigated mRNA abundance of ACE2 and associated SARS-CoV-2 cell machinery that can influence ACE2 (ADAM17, TMPRSS2) in lung and adipose tissue from low density lipoprotein receptor (*Ldlr*) deficient male and female mice fed a Western diet to mimic other risk factors for COVID-19 severity (obesity, hypertension). We focused on lung and adipose tissue because the lung serves as the major entry site for SARS-CoV-2. Adipose tissue was a focus because this tissue has a complete RAS, including ACE2(21, 22), and obesity is a risk factor for COVID-19 severity(23–26). We then defined effects of SARS-CoV-2 S protein on ACE2 activity in alveolar type II (ATII) lung epithelial cells and 3T3-L1 adipocytes. Next, we investigated effects of administration of SARS-CoV-2 S protein on the lung and adipose RAS of male and female *Ldlr^-/-^* mice. Finally, we quantified ACE2 activity and AngII content in lung specimens from a bilateral lung transplant with significant fibrosis for a patient with confirmed COVID-19 and a fibrotic lung control.

## Materials and Methods

### Primary isolates of lung ATII cells

Alveolar type II (ATII) cells were isolated from male mice as described previously.(27) Briefly, lungs were removed and digested in dispase, the resulting cell suspension was filtered, and ATII cells were separated by differential adherence to culture plates coated with exclusion antibodies.(28, 29). Cells were then gently panned from the plates, collected and centrifuged. Cells were resuspended in mouse ATII growth medium and cultured on fibronectin-coated cell culture plates. On day 2, media was changed and cells were used for experiments on day 3.

### 3T3-L1 adipocytes

Murine 3T3-L1 cells were obtained from ATCC (CL-173) and maintained in DMEM containing fetal bovine serum (FBS, 10%). Cells were differentiated to adipocytes in DMEM (10% FBS), human insulin (0.2 μM; Novolin R 100 U/ml, ReliOn), 3-isobutyl-1-methylxanthine (0.5 mM, Catalog I6879, Sigma-Aldrich), dexamethasone (1.0 μM, Catalog# 194561, MP Bio), and rosiglitazone (2 μM; Catolog# 194561, Sigma-Aldrich) for three days. Cell were then transferred to maintenance media (DMEM, 10% FBS, 0.2 μM insulin) for a total of 8 days. For measurements of ACE2 activity, washed differentiated adipocytes (triplicates) were incubated in Krebs buffer (1.5 mls, 37°C, 5% CO_2_) containing the angiotensin converting enzyme inhibitor enalapril (20 μM; Catalog# PHR1289, Sigma-Aldrich) to eliminate further conversion of angiotensin I to AngII, Z-Pro-prolinal (200 μM, Catalog# SML0205, Sigma-Aldrich) to inhibit prolyloligopeptidase(30), or MLN-4760 (200 μM;(31)) to inhibit ACE2. Following addition of the ACE2 substrate, buffer was removed from incubated cells at specified times (0, 1, 2, 3, 4, 5, 15, 16 hours) and stored at 4°C for quantification of ACE2 activity as described below. In some studies, conditioned media was removed from cells at specified time points following incubation with SARS-CoV-2 S protein prior to addition of the ACE2 substrate and quantification of ACE2 activity (sACE2).

To define effects of SARS-CoV-2 S protein on adipocyte ACE2 activity, differentiated adipocytes were incubated with vehicle or SARS-CoV-2 S protein (33 nM) for 24 hours (37°C, 5% CO_2_). Buffer was removed from cells to quantify sACE2 using the ACE2 activity assay as described below. Washed adipocytes were used to quantify cACE2 activity.

### ACE2 activity

ACE2 activity was quantified in cultured cells or adipose tissue explants.(17, 32) Samples were incubated in buffer (75 mM Tris-HCl, pH 7.5; 200 mM NaCl, 10 μM ZnCl) plus a series of inhibitors (enalapril, 20 μM; Z-Pro-Prolinal 100 μM; with or without MLN-4760 (20 μM). These inhibitors were used to prevent further production of AngII (enalapril), inhibit prolyoligopeptidase (Z-Pro-Prolinal), or inhibit ACE2 (MLN-4760). An ACE2 fluorogenic substrate, Mca-APK(DPN) (50 μM; Enzo Catalog# BML-P163-0001) was incubated with cells or explants at 37°C, 5% CO2 and then samples were placed in a fluorescence plate reader (excitation, 320 nm; emission, 405 nm). ACE2 activity (RFU/hr/μg protein) was calculated for each condition, and specific ACE2 activity was defined as that in the presence of Z-Pro-Prolinal minus residual activity present in the presence of MLN-4760.

### AngII quantification

A small amount of frozen lung (≈50 mg) was homogenized (GenoGrinder, 1,350 rpm, 1 minute) in buffer (potassium phosphate buffer, 10mm, pH 7.4; EDTA, 3 mM; 0.15 mM 8-hydroxyquinoline) at 4°C. The homogenate was centrifuged (14,000 rpm, 10 minutes, 4°C), and supernatants were transferred to fresh tubes. RIA was performed using a rabbit-anti-angiotensin II antiserum (0.3 μg/ml; Peninsula Laboratories Catalog# T-4005) and [^125^I]AngII (2,175 Ci/mmol; ≈12,000 cpm/ml; Georgetown University) as tracer with samples incubated for 24 hours at 4°C. AngII standards (10 – 2,500 pg/ml) were used to generate a standard curve which was used to quantify lung AngII. Antibody bound AngII was separated from free AngII by adsorption to charcoal (0.5%, Fisher Cat C170). Radioactivity (bound fraction) was quantified by gamma counting. AngII levels were normalized to lung protein content (BCA protein assay; Pierce Catalog# 23227).

### Animals

To define sex differences in mRNA abundance of ACE2 and associated cell machinery, male and female *Ldlr^-/-^* mice (2 months of age; n = 4 mice/group/sex) were obtained from the Jackson Laboratory. Mice of the Four Core Genotypes (FCG), an original gift from Dr. Art Arnold (UCLA), were bred onto an *Ldlr^-/-^* background as defined previously.(33) This model has a mutation in *Sry,* the testes determining gene of the Y chromosome, which was inserted onto autosomes of male mice to enable fertility.(34, 35) Separation of *Sry*-dependent testes formation from the presence of the Y chromosome allows for creation of the FCG. When XY^Sry^ FCG male *Ldlr^-/-^* mice are bred to XX females, FCG are generated: XX female (XXF), XY^-^F, XY^Sry^ male (XYM) and XX^Sry^ male (XXM). Mice of these genotypes were either sham-operated or gonadectomized (GDX) at 2 months of age under anesthesia (ketamine/xylazine, 100/10 mg/kg, ip) using previously established methods(36). Two weeks later, all mice were fed a Western diet (TD.88137; Teklad) for a period of 4 months to mimic risk factors of COVID-19 severity (obesity, hypertension). At study endpoint, tissues (lung, gonadal adipose tissue) were removed from euthanatized mice.

To investigate effects of SARS-CoV-2 S protein administration on tissue RAS, *Ldlr^-/-^* male and female mice (2 months of age, n = 4 mice/group/sex; the Jackson Laboratories) were fed the Western diet for one week. Mice were administered vehicle (saline) or full length recombinant SARS-CoV-2 S protein (T4 foldon trimerization domain to form a protein trimer, which approximates the structure on the virus; Protein Core Service Facility, University of Kentucky, molecular weight of ~180,000 validated by SDS-PAGE). The dose (2 nmol/kg of the SARS-CoV-2 S protein trimer) was based on previous studies administering recombinant SARS-CoV S protein to mice.(37) Male and female mice were administered 3 divided doses (2 nnmol/kg, ip) separated by 1 hour. Tissues (lung, gonadal adipose tissue) were removed from anesthetized mice at study endpoint.

All animal protocols were reviewed and approved by the University of Kentucky Institutional Animal Care and Use Committee and adhere to the guidelines of the National Institutes of Health.

### Human Lung

Human lungs were obtained under the IRB approved protocol for discarded human tissues. Samples were obtained from patients undergoing bilateral lung transplant at University of Kentucky Health Care. Lungs were dissected into small pieces that were snap frozen in threaded cryovials in liquid nitrogen until use.

ACE2 activity in human lung explants was measured by cutting small pieces (explants, ~20 mg) of lung tissue and placing the pieces in the wells of a 48-well plate containing 0.5 mL of Krebs Buffer containing enalapril (10 uM) and Z-pro-prolinal (200 uM). Tissues were then incubated at 37°C for 45 minutes. The buffer was removed and replaced with 0.5 mL fresh Krebs Buffer with inhibitors plus the ACE2 substrate Mca-APK-Dpn (30 uM) and incubated at 37°C. Sixty uL samples were removed from each well at 0, 1, 2, 3, and 5 hours after the addition of substrate. The samples were stored frozen until fluorescence could be measured using Ex = 320 nm, Em = 405 nm.

AngII activity in lung tissue was measured by RIA. Briefly, lung tissue was homogenized in 10 mM potassium phosphate buffer, pH 7.4, 3 mM EDTA, 0.15 mM 8-hydroxyquinoline, centrifuged at 16,000 x g for 10 minutes at 4°C. The supernatants were then measured for AngII by radioimmunoassay (RIA) with a competitive assay using [^125^I] labeled AngII as described above.

### Statistics

All data are presented as means ± SE. Data were analyzed and graphed using GraphPad Prism 9.0.2 software. Group comparisons were performed using a two-way ANOVA or unpaired t-test where appropriate. For studies comparing male and female mice, a two-way ANOVA with sex and treatment (vehicle, SARS-CoV-2 S protein) as between group factors and Tukey’s post hoc test. Statistical significance was accepted with P<0.05.

## Results

### Sex hormones and an XX sex chromosome complement regulate lung ACE2 mRNA abundance in Western diet-fed *Ldlr^-/-^* mice

ACE2 gene is on the X chromosome, and studies indicate that sex hormones, primarily estrogen, positively regulate ACE2 gene expression in various tissues.(15–20) We used male and female FCG mice, with and without GDX to define effects of sex hormones and fed a Western diet to simulate COVID-19 risk factors (obesity, hypertension). FCG mice enabled determination of the relative effects of sex hormones *versus* sex chromosome complement on lung ACE2 mRNA abundance. As shown in Figure 1A, ACE2 lung mRNA abundance was not different between intact male and female mice (P>0.05), with significant reductions in ACE2 mRNA abundance in lungs of GDX male and female mice (P<0.05). To define a role for sex chromosomes, we compared ACE2 mRNA abundance in lungs of XXF, XXM, XYM and XYF FCG mice. Mice with an XX sex chromosome complement, whether they were gonadal males or females, had higher lung ACE2 mRNA abundance than XY FCG mice (Figure 1B; P<0.05). Since sex hormones and an XX sex chromosome complement both influenced lung ACE2 mRNA abundance, we examined interactions between these sex defining factors in XXF and XYF intact and GDX mice. Interestingly, lung ACE2 mRNA abundance was decreased by GDX in XXF, but not XYF FCG mice (Figure 1C; P<0.05).

**Figure 1.**
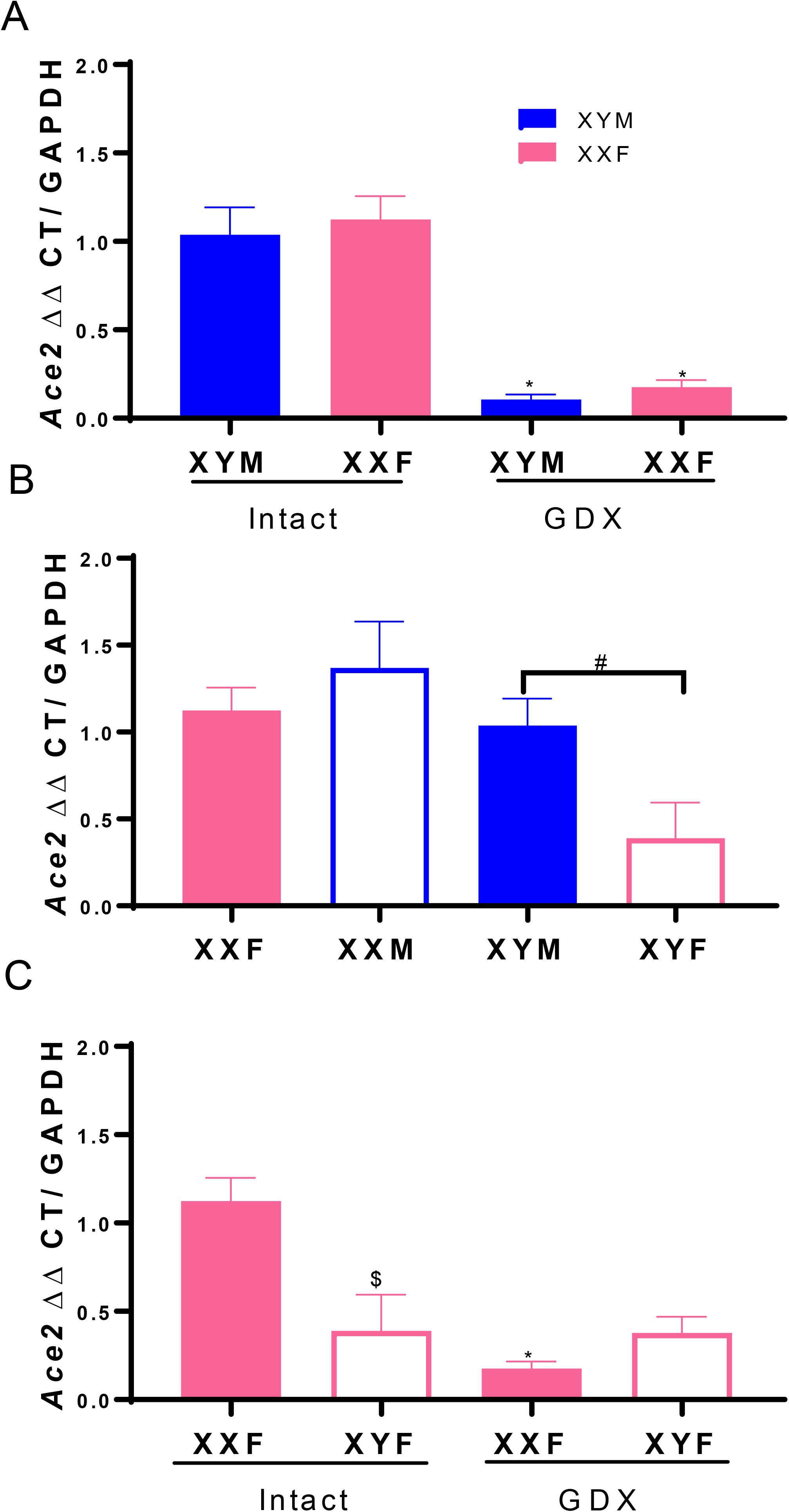
Sex hormones and an XX sex chromosome complement regulate lung ACE2 mRNA abundance. A, ACE2 mRNA abundance in lung from intact and gonadectomized (GDX) XY male (XYM) and XX female (XXF) mice. B, ACE2 mRNA abundance in lungs from FCG mice: XXF and XYM mice, solid bars; XX male (XXM) and XY female (XYF) mice, open bars. C, ACE2 mRNA abundance in lungs from FCG female (XXF, XYF) intact or GDX mice. Data are mean ± SEM from n = 3-4 mice/group. Analyses performed using two-way ANOVA with pairwise comparisons. A: *, P<0.05 compared to intact within sex chromosome complement. B: #, P<0.05 compared to XX. C. *, P<0.05 compared to intact within sex chromosome complement. $, P<0.05 compared to XXF within intact mice.

### Male sex hormones regulate lung mRNA abundance of ADAM17 and TMPRSS2

Lungs from intact male mice had higher mRNA abundance of ADAM17 than intact females (Figure 2A; P<0.05). Moreover, GDX significantly decreased lung ADAM17 mRNA abundance in male but not female mice (Figure 2A; P<0.05). In contrast, lung mRNA abundance of TMPRSS2 was not different between male and female mice (Figure 2B; P>0.05). However, following GDX, TMPRSS2 mRNA abundance decreased in lungs of male, but not female mice (Figure 2B; P<0.05).

**Figure 2.**
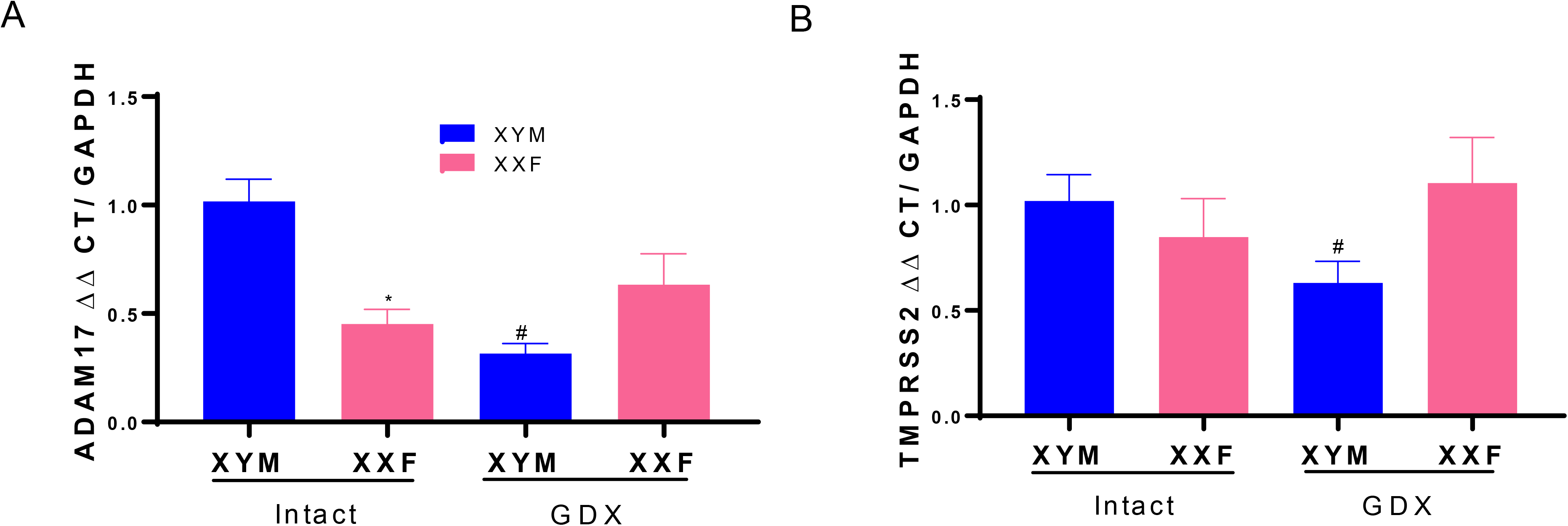
ADAM17 and TMPRSS2 mRNA abundance in lung are regulated by male sex hormones. A, ADAM17 mRNA abundance in lungs of intact and gonadectomized (GDX) male and female mice. B, TMPRSS2 mRNA abundance in lungs of intact and GDX male and female mice. Data are mean ± n = 4 mice/group. Analyses performed using two-way ANOVA with pairwise comparisons. *, P<0.05 compared to male within intact. #, P<0.05 compared to intact within sex chromosome complement.

### Female sex hormones and an XX sex chromosome complement stimulate adipose ACE2 mRNA abundance

We demonstrated previously that female sex hormones positively regulate adipose ACE2 mRNA abundance through an estrogen, estrogen receptor-α (Erα)-mediated mechanism at the ACE2 gene promoter.(16) In this study, we used FCG mice to contrast effects of sex hormones and sex chromosome complement on adipose ACE2 mRNA abundance. In adipose tissue of Western diet-fed *Ldlr^-/-^* male and female FCG mice, ACE2 mRNA abundance was higher in XXF than XYM mice (Figure 3A; P<0.05). In addition, adipose ACE2 mRNA abundance was higher in XX than XY mice regardless of gonadal sex (Figure 3A; P<0.05).

**Figure 3.**
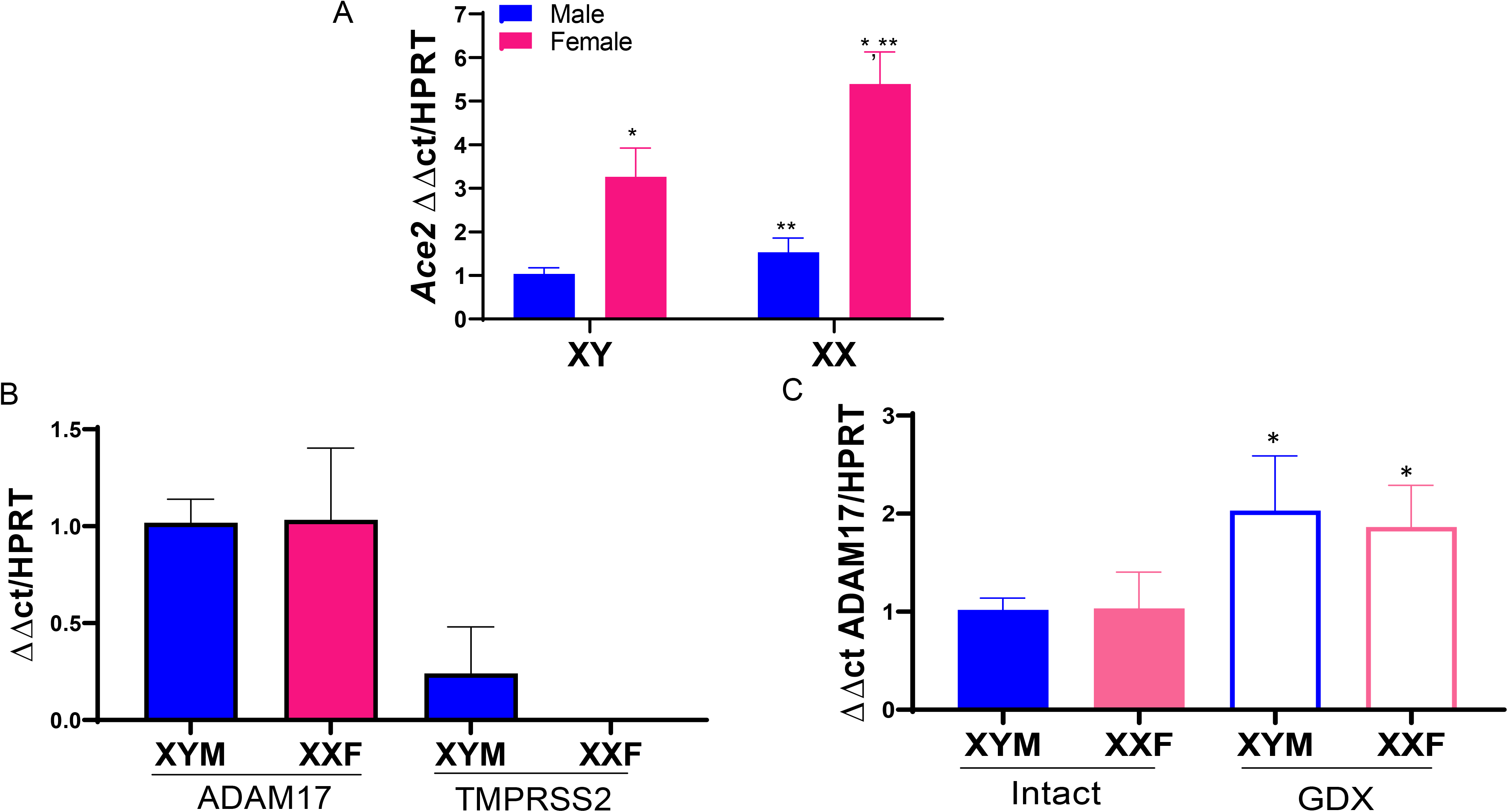
ACE2 mRNA abundance in adipose is regulated by sex hormones and sex chromosomes, while ADAM17 mRNA abundance is regulated by sex hormones in adipose tissue. A, ACE2 mRNA abundance in adipose tissue of FCG XX female (XXF), XY female (XYF), XY male (XYM) and XX male (XXM) mice. B, Relative mRNA abundance of ADAM17 and TMPRSS2 in adipose tissue of male and female mice. C. Effect of GDX on ADAM17 mRNA abundance in adipose tissue. Data are mean ± SEM from n = 4 mice/group. *, P<0.05 compared to male within sex chromosome genotype. **, P<0.05 compared to XYF.

### Adipose mRNA abundance of ADAM17 is regulated by sex hormones

Adipose tissue is known to express ADAM17 which serves as TNF-α converting enzyme and also is capable of shedding the ectodomain of ACE2.(11, 12, 21) In contrast, adipose tissue has not been extensively studied in the context of SARS-CoV-2 infectivity, which would require machinery such as TMPRSS2 to cleave the S proteins of SARS-CoV-2.(38, 39) We contrasted mRNA abundance of ADAM17 to TMPRSS2 in adipose tissue from male and female mice. ADAM17 mRNA abundance in adipose tissue did not differ between male and female mice, with expression levels higher than TMPRSS2 mRNA abundance which was only detected in adipose tissue of male mice (Figure 3B). ADAM17 mRNA abundance was higher in adipose tissue of GDX male and female mice compared to same sex intact mice (Figure 3C; P<0.05).

### SARS-CoV-2 S protein reduces ATII and 3T3-L1 adipocyte ACE2 activity

Previous investigators have demonstrated that SARS-CoV-1 S protein reduced cell surface levels of ACE2.(6–8, 37) We defined effects of SARS-CoV-2 S protein on ACE2 activity in lung ATII cells and differentiated 3T3-L1 adipocytes. Incubation of SARS-CoV-2 S protein with lung ATII cells from male mice resulted in a dose-dependent reduction in cACE2 activity (Figure 4A). In differentiated 3T3-L1 adipocytes incubated with SARS-CoV-2 S protein, we quantified ACE2 activity when the ACE2 substrate was incubated with intact cells (cACE2) and when the ACE2 substrate was added to conditioned media removed from adipocytes (sACE2) after exposures to S protein. SARS-CoV-2 S protein significantly decreased cACE2 activity in adipocytes (Figure 4B; P<0.05) while resulting in a modest, but not significant increase in sACE2 activity in conditioned media (Figure 4B; P>0.05).

**Figure 4.**
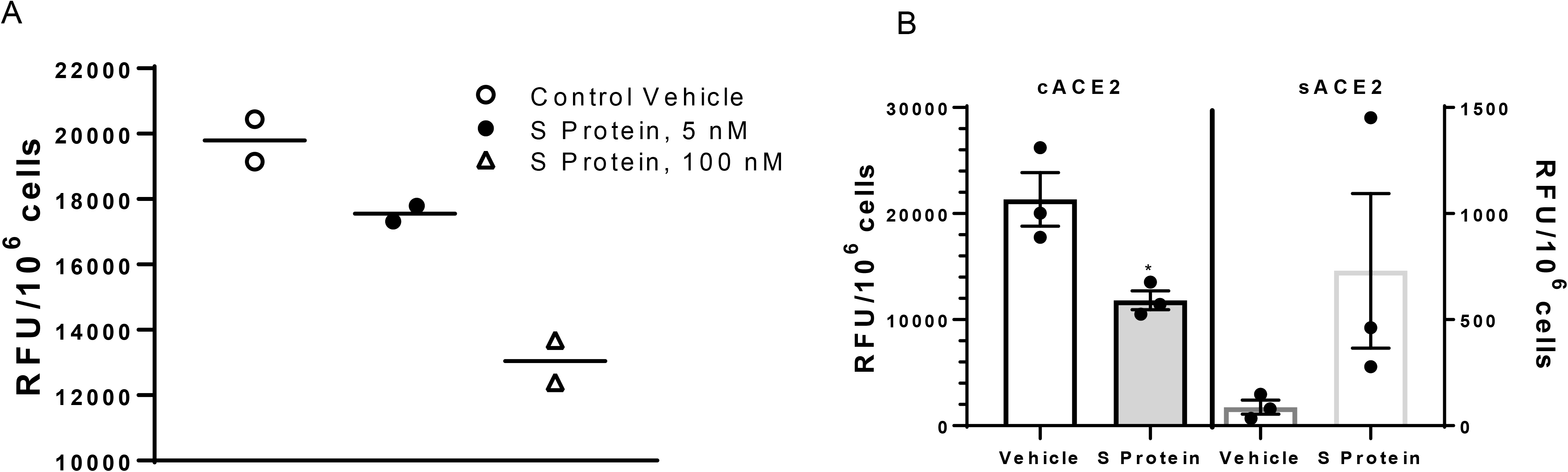
SARS-CoV-2 S protein decreases ACE2 activity in lung ATII cells and 3T3-L1 adipocytes. A, SARS-CoV-2 dose-dependently reduces ACE2 activity in lung ATII cells. B, SARS-CoV-2 S protein (100 nM) reduces cell ACE2 (cACE2) activity while increasing soluble ACE2 (sACE2) activity in conditioned media from incubated adipocytes. A, Data are triplicates from n = 2 primary isolates. B, Data are mean ± SEM from n = 3 replicates. *, P<0.05 compared to vehicle.

### SARS-CoV-2 S protein administration increases lung AngII content and decreases adipose ACE2 activity to a greater extent in male than female mice

Previous investigators demonstrated that administration of SARS-CoV-1 S protein to mice decreased lung ACE2 protein content and elevated AngII levels in lung.(37) However, the sex of the mice used in these studies was not identified. Using a similar dosing protocol as described previously(37), we administered SARS-CoV-2 S protein to male and female *Ldlr^-/-^* mice fed a Western diet for one week. Lung AngII content was significantly increased by SARS-CoV-2 S protein compared to vehicle controls in male, but not female mice (Figure 5A; P<0.05). Adipose ACE2 activity was higher in male than female adipose tissue of vehicle-injected mice (Figure 5B; P<0.05). Moreover, administration of SARS-CoV-2 S protein resulted in a significant reduction of adipose ACE2 activity in male, but not female mice (Figure 5B; P<0.05).

**Figure 5.**
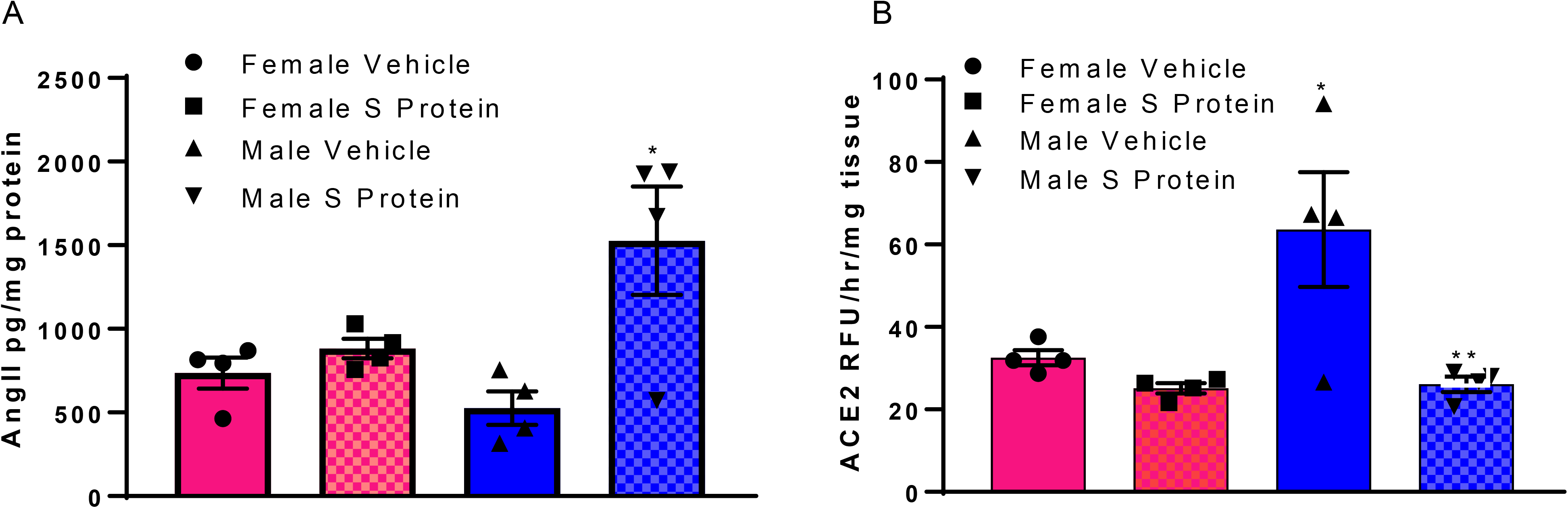
SARS-CoV-2 S protein administration increases lung AngII content and decreases adipose ACE2 activity in male but not female mice. A, Administration of SARS-CoV-2 Spike protein (S protein; 2 nmol/kg in three divided doses, i.p.) results in an increase in AngII content in lungs of male but not female mice. B, SARS-CoV-2 S protein decreases adipose ACE2 activity in male but not female mice. Data are mean ± n = 4 mice/group/sex. *, P<0.05 compared to female within treatment. **, P<0.05 compared to vehicle within sex.

### Human lung from a confirmed COVID-19 patient has reduced ACE2 activity and elevated AngII content

We obtained a specimen of human lung from a patient undergoing bilateral lung transport due to significant resultant pulmonary fibrosis from a confirmed diagnosis of COVID-19 and compared ACE2 activity and AngII content to that of a lung specimen from a patient with lung fibrosis. ACE2 activity and lung AngII content were quantified in three separate samples from each patient. ACE2 activity was lower in COVID-19 lung compared to fibrotic lung control (COVID-19: 270 ± 31; Fibrotic Control: 470 ± 45 RFU/mg/hr; P<0.05) while AngII content was higher in COVID-19 lung compared to fibrotic lung control (COVID-19: 206 ± 23; Fibrotic Control: 124 ± 20 pg/mg tissue).

## Discussion

In this study we report that sex hormones and sex chromosome complement influence critical SARS-CoV-2 machinery, with a focus on ACE2, in pivotal target tissues implicated in COVID-19 severity. Sex-dependent regulation of ACE2 favored higher residual cACE2 and greater local control of the RAS in lung and adipose from females compared to males exposed to SARS-CoV-2 S protein. Moreover, enzymes capable of shedding the ectodomain of cACE2 were regulated by sex hormones in a tissue-specific manner that also favored activation of the local RAS in males compared to females exposed to SARS-CoV-2 S protein. Exposure of adipocytes and ATII cells to SARS-CoV-2 S protein decreased ACE2 activity. Administration of SARS-CoV-2 S protein to male, but not female mice, increased lung AngII content and reduced adipose ACE2 activity. Finally, human lung specimens from a male patient with confirmed COVID-19 had lower ACE2 activity and higher AngII content than that from a male patient with fibrotic lung. Taken together, these results suggest that sex-dependent regulation of ACE2 and other SARS-CoV-2 machinery in pivotal cell types may contribute to an activated local RAS and augmented severity of COVID-19 in males.

A growing body of literature suggests that while infection rates of males and females are similar for COVID-19, the severity of the condition is worse in men than women, culminating in worse prognosis and greater death rates of males. ACE2 is the major receptor for entry of SARS-Co-V-1 and 2 into cells.(4, 5 2020 2020) Thus, it is important to understand factors controlling ACE2 expression in cells, and whether these factors, such as sex-defining influences, influence infection rates and severity of the condition. In the few studies performed to date, levels of ACE2 have not been conclusively linked to infection rates amongst affected individuals.(40, 41) Thus, regulatory mechanisms for control of ACE2 expression do not appear to have an immense impact on SARS-CoV-2 infection rates. Early studies with SARS-CoV-1 demonstrated that viral binding to ACE2 resulted in subsequent internalization of the protein via clathrin-mediated endocytosis that leads to a loss of functional ACE2 in the cell membrane.(8, 37) Associated with loss of cACE2, lung AngII content rose and acute lung injury was augmented following exposure to SARS-CoV-1 (full length virus or S protein) which could be ameliorated by administration of an angiotensin type 1 receptor (AT1R) antagonist.(37) These results suggested that rather than infection rates, cell surface levels of ACE2 may impact local control of the RAS and COVID-19 severity. Our results demonstrate similar levels of ACE2 mRNA abundance in lungs of age-matched and Western diet-fed male and female mice, supportive of a similar rate of SARS-CoV-2 infection between sexes as defined previously.(42) Notably, following GDX of males and females, lung ACE2 mRNA abundance declined markedly, suggesting that reductions in sex hormone levels in the aging population and lower residual levels of ACE2 following SARS-CoV-2 exposure may contribute to higher COVID-19 severity in aging males and females. However, since COVID-19 infection rates also increase with age(43), declines in ACE2 mRNA abundance in GDX males and females, similar to andropause and menopause within this population, are not consistent with higher COVID-19 infection rates of this population. The finding of lower ACE2 activity associated with higher AngII content in lung specimens from a COVID-19 patient compared to fibrotic lung control tissue supports further studies addressing the hypothesis that loss of cell surface ACE2 following SARS-CoV-2 exposure impacts the local RAS.

We found that an XX sex chromosome complement contributed to higher ACE2 expression in lungs and adipose tissue of females than males. ACE2 is on the X chromosome, but it is unclear if ACE2 escapes the process of XCI which would result in higher ACE2 expression levels in females than males. Our results support escape from XCI of ACE2 in lung and adipose tissue, that coupled with influences of sex hormones on ACE2 expression levels, could influence the residual level of ACE2 following SARS-CoV-2-mediated ACE2 internalization. In support of this suggestion, male mice exposed to SARS-CoV-2 S protein exhibited higher lung AngII content and lower adipose ACE2 activity than females, supporting a more activated local RAS in males than females that might influence COVID-19 severity.

Moreover, our results demonstrate interactions between sex hormones (estrogen) and sex chromosomes (XX) to stimulate ACE2 expression in lung. Namely, estrogen-mediated stimulation of lung ACE2 gene expression required an XX sex chromosome complement, which may have ramifications for transgender therapy.

In addition to lung, we defined ACE2 expression levels and response to SARS-CoV-2 S protein in adipose tissue of male and female mice. We focused on adipose tissue since obesity is a risk factor for COVID-19 severity(23–26), and since adipocytes possess a complete RAS including ACE2(21). Moreover, previous studies in our laboratory demonstrated that adipose tissue becomes a significant source of systemic AngII in the development of obesity-hypertension in male mice.(44) In addition, deficiency of ACE2 in adipocytes promoted the development of obesity-hypertension in female, but not male mice.(45) Thus, SARS-CoV-2 regulation of adipose ACE2 may have systemic influences on the RAS and cardiovascular control in a sex-specific manner, especially in obese subjects, thereby influencing COVID-19 severity. In contrast to lung, but in agreement with previous findings(21), adipose ACE2 mRNA abundance was higher in females than males. Notably, an XX sex chromosome complement resulted in higher ACE2 mRNA abundance in adipose tissue, similar to lung, supporting escape of ACE2 from XCI. These findings support sex differences in adipose ACE2 that may relate to COVID-19, but it is unclear if SARS-CoV-2 circulates to infect or influence adipocytes. Our results demonstrate that some proteins of the SARS-CoV-2 machinery (ACE2, ADAM17) are expressed in adipose tissue, while other important proteins involved in viral cell entry (TMPRSS2) are expressed at very low levels, indicating that infection of adipocytes may not be extensive. However, given that exposures to SARS-CoV-2 S protein decreased ACE2 activity *in vitro* and *in vivo,* it is possible that if the virus circulated to adipocytes infection and regulation of the adipose RAS may occur. In support of this, measurements of cell-free DNA as a biomarker of injury in patients with COVID-19 demonstrated profiles indicative of adipose tissue and associated with end-organ damage in subjects with worse COVID-19 outcomes.(46)

In addition to ACE2, we defined effects of sex-defining factors on expression of ADAM17 and TMPRSS2 in lung and adipose tissue. These enzymes, implicated in viral entry, also shed the ectodomain of cACE2 and thereby activate the local RAS. There is limited information on regulation of ADAM17 by sex hormones and/or sex chromosomes. Our results indicate that ADAM17 is positively regulated by male sex hormones with no influence of female sex hormones in lung, while both sex hormones negatively regulate ADAM17 expression levels in adipose tissue. In lung, these results would suggest higher shedding of cACE2 and activation of the local RAS in males, while influences of sex differences in ADAM17 expression in adipose tissue are more difficult to predict. TMPRSS2 is known to exhibit strong androgen receptor promoter regulation and has been suggested as a mechanism contributing to sex differences in COVID-19 severity.(47) Our results support this hypothesis, as removal of male sex hormones reduced TMPRSS2 mRNA abundance in lung and adipose tissue of male mice. In adipocytes exposed to SARS-CoV-2 S protein, cACE2 activity declined while detectable activity of sACE2 in conditioned media from adipocytes rose, supporting SARS-CoV-2 S protein-mediated shedding of cACE2. Shedding of the ectodomain of cACE2 containing the viral RBD would be predicted to reduce infectivity of SARS-CoV-2 at adipocytes. Indeed, an ongoing clinical trial utilizes recombinant sACE2 to bind SARS-CoV-2 and reduce its ability to interact with cells through cACE2.(48–50) Our interest is the consequence of SARS-CoV-2 S protein induced shedding of cACE2 on the adipose RAS, especially in the context of obesity where adipose-derived AngII has been demonstrated to promote obesity-induced hypertension of male mice.(44) In support, administration of SARS-CoV-2 S protein to Western diet-fed male, but not female mice reduced adipose ACE2 activity, which would predictably increase adipose-derived levels of AngII. Future studies will determine the pathophysiological impact of SARS-CoV-2 S protein-mediated regulation of adipose ACE2 in the context of obesity.

In conclusion, results from this study suggest that sex defining factors, including sex hormones and sex chromosome complement, influence expression of ACE2 and other SARS-CoV-2 machinery (ADAM17, TMPRSS2) to favor a higher residual level of cACE2 and lower local RAS activity in females than males exposed to SARS-CoV-2 S protein. Reduced lung ACE2 activity and higher AngII content in a patient with COVID-19 compared to fibrotic lung disease support the implications of loss of cell surface ACE2 on the local lung RAS. Future studies should define the impact of these findings of sex-dependent regulation of ACE2 and its sheddases in pivotal target tissues on disease severity and whether these effects persist in subjects with post-acute sequalae of SARS-CoV-2 infection.

## Acknowledgments

We thank Ms. Victoria English for her assistance with animal handling and laboratory management and Ms. Heba Ali for technical assistance in various aspects of the study design. We thank the University of Kentucky COVID-19 biobank for acquisition of study specimens and/or data.

